# Extensive Benchmarking of Community Detection Algorithms

**DOI:** 10.1101/2025.05.07.652778

**Authors:** R Sapna, Harikeshav Karthik, Karthik Raman

## Abstract

The detection of clusters or community in networks is an important problem in network science. We systematically evaluate many widely used community detection algorithms and their variants to identify clusters in complex networks. As the ground truth for assessing accuracy, we use artificial networks modeled on power-law distributions and real-world social networks. In addition, we also performed gene enrichment analysis on human and yeast protein–protein interaction networks to evaluate algorithms on their ability to uncover enriched communities. We implement and adapt an extensive suite of classical algorithms and their modern variants, classified into five types: stochastic, kernel-based, modularity-based, hierarchical, and local search-based. The algorithms are benchmarked primarily using the Normalized Mutual Information metric, with additional analyses focused on granularity by examining cluster ratio and computational time complexity. We find that decreasing the modularity of networks leads to a consistent decline in performance that follows a sigmoidal trajectory as communities become less defined. Algorithms with greater granularity remain stable when community structures are less distinct, while computation time remains independent of network modularity. Additionally, algorithms tend to perform poorly on smaller networks, and higher accuracy often requires a time complexity trade-off for specific high-performing methods. However, as the analysis expands to more extensive networks, this trade-off becomes more pronounced, highlighting the need for efficient scalability. Based on our benchmark and gene enrichment analysis results, we also present recommendations to practitioners. Our robust Python package, complete with a user-friendly command-line interface, empowers users to easily apply these algorithms to their datasets.

**Author summary:** In the era of big data, clustering has become an essential tool for processing and analyzing vast amounts of information. By dividing large data sets into smaller, meaningful clusters, we can simplify complex data structures, parallelize tasks, and reveal hidden patterns. This has become an essential preprocessing step that is widely used in various computational domains. Although traditional clustering algorithms remain popular, our work implements range of hybrid techniques that combine classical methods and some novel approaches. We also benchmark the performance of these approaches, their efficiency and efficacy in handling different types of networks.

## 1 Introduction

Community detection algorithms are crucial tools for identifying clusters or groups of densely connected nodes within networks [1–5]. These algorithms have wide-ranging applications [6], from sociology [7–9] and biology [10–12] to big data processing [13–15] and parallelizing computational tasks [16]. They provide valuable insights into the structure and organization of complex systems by uncovering hidden patterns [17] and mesoscopic properties [18] of the underlying network.

Historically, benchmarking efforts have focused on using standard methods available in Python and R libraries such as igraph and NetworkX [19, 20]. A significant aspect of the present work comes from the DREAM Challenge (2018) [21], which focused on identifying disease-enriched modules in extensive anonymized biological networks.

More recent developments include techniques that employ deep learning techniques [22–24] and overlapping community detection algorithms [25, 26], both of which have gained considerable traction.

We implement hybrid techniques that enhance traditional algorithms through various modifications and introduce novel approaches. We create a comprehensive framework through extensive benchmarking to evaluate and compare the effectiveness of different community detection methods in diverse network structures. We also provide the algorithms implemented as a Python package, making them an accessible tool for future research and applications.

## 2 Results

In this section, we compare various community detection algorithms based on their performance. The overall benchmarking workflow is illustrated in Fig. 1.

**Fig 1.**
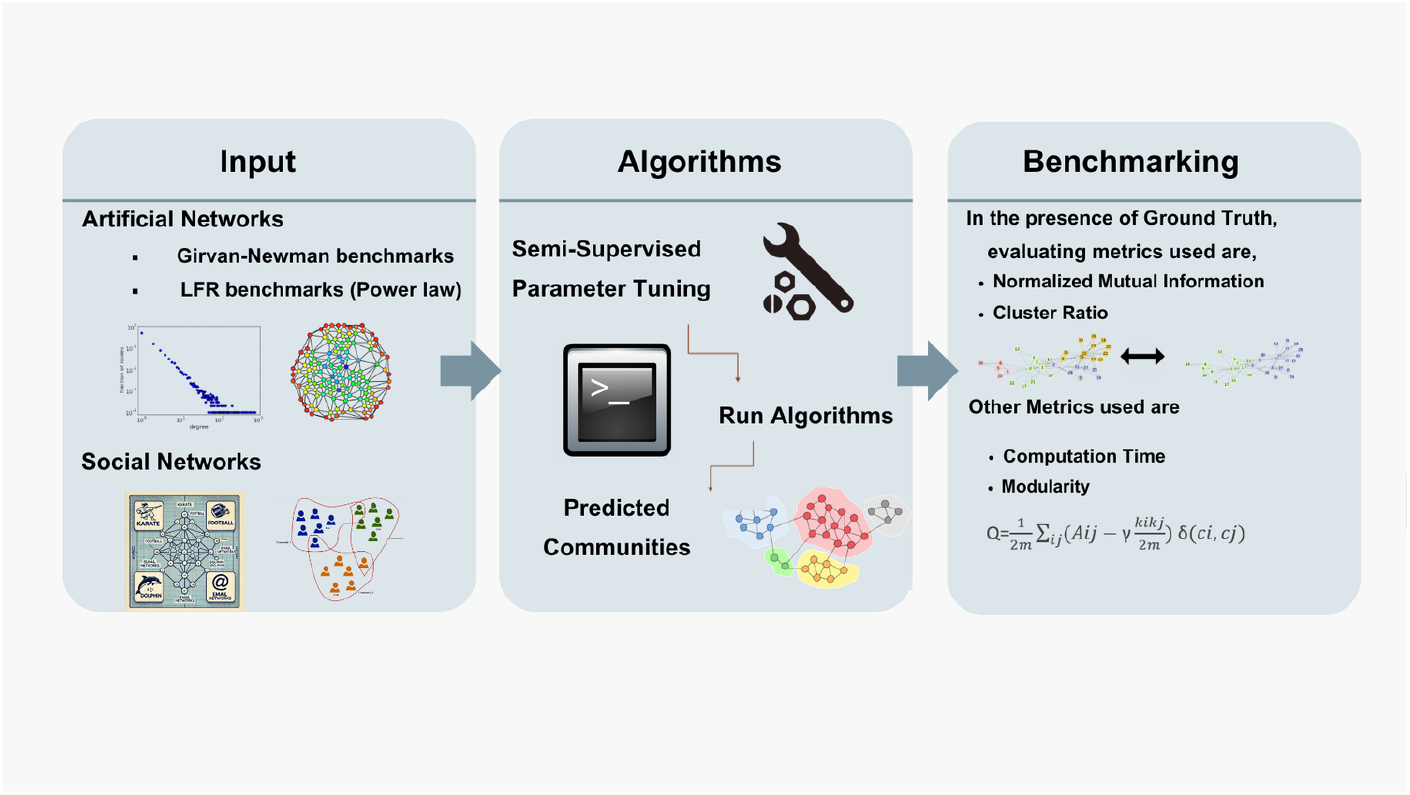
Overall Benchmarking Workflow. The benchmarking workflow can be organized into three stages as depicted above.

### 2.1 Artificial Benchmarks: Mixing parameter *µ* based analysis

In the case of artificial benchmark analysis, we chose the weighted and undirected Lancichinetti–Fortunato–Radicchi (LFR) networks following power-law distributions for this study. The details of the parameters used to generate the networks are given in Table 3. Two types of benchmark analysis have been performed using the LFR networks shown in Fig. 2,Fig. 3,Fig. 4,Fig. 5,Fig. 6 and Fig. 7.

**Fig 2.**
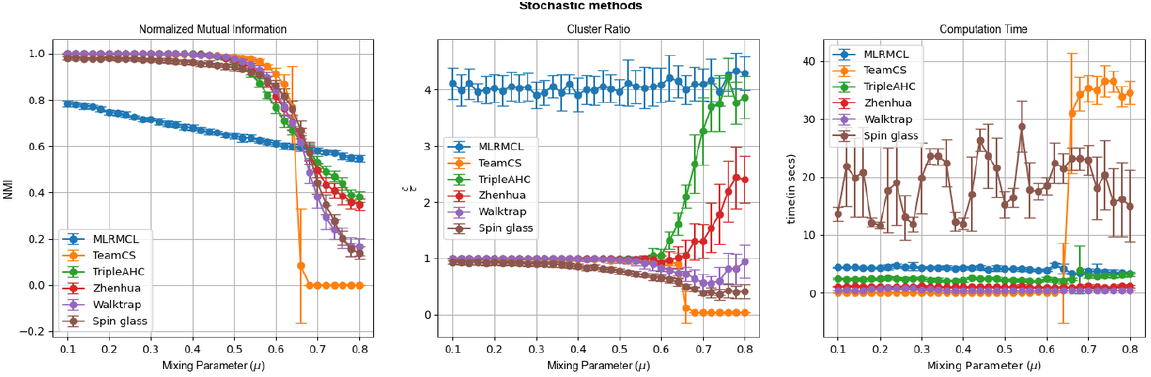
Mixing parameter(*µ*) based analysis on Stochastic methods. The *µ* of the LFR network is varied from 0.1 to 0.8.

**Fig 3.**
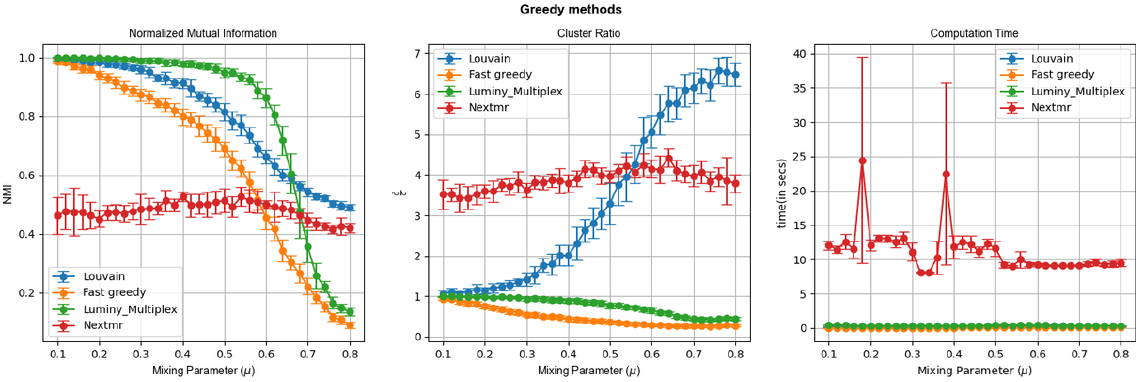
Mixing parameter(*µ*) based analysis on Greedy methods. The *µ* of the LFR network is varied from 0.1 to 0.8.

**Fig 4.**
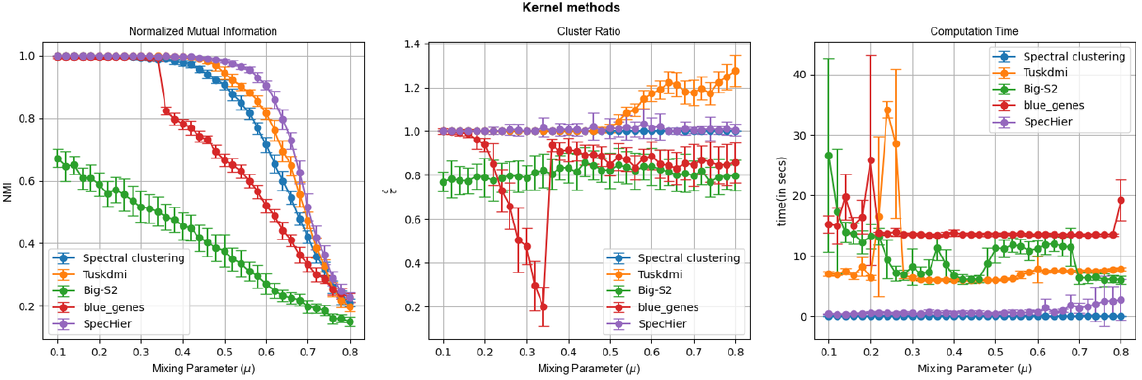
Mixing parameter(*µ*) based analysis on Kernel methods. The *µ* of the LFR network is varied from 0.1 to 0.8.

**Fig 5.**
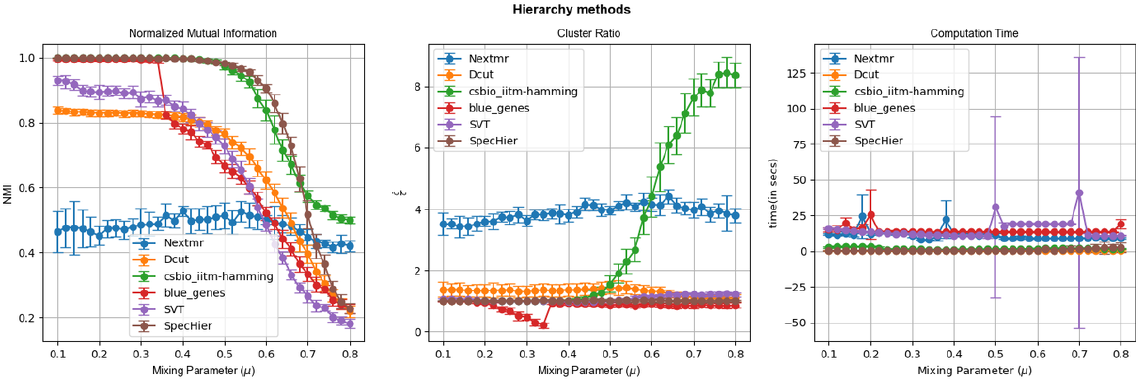
Mixing parameter(*µ*) based analysis on Hierarchy methods. The *µ* of the LFR network is varied from 0.1 to 0.8.

**Fig 6.**
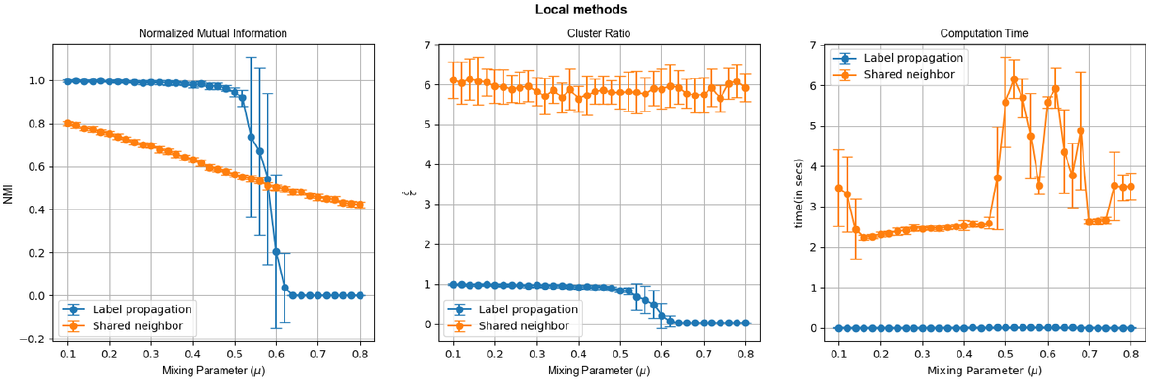
Mixing parameter(*µ*) based analysis on Local methods. The *µ* of the LFR network is varied from 0.1 to 0.8.

**Fig 7.**
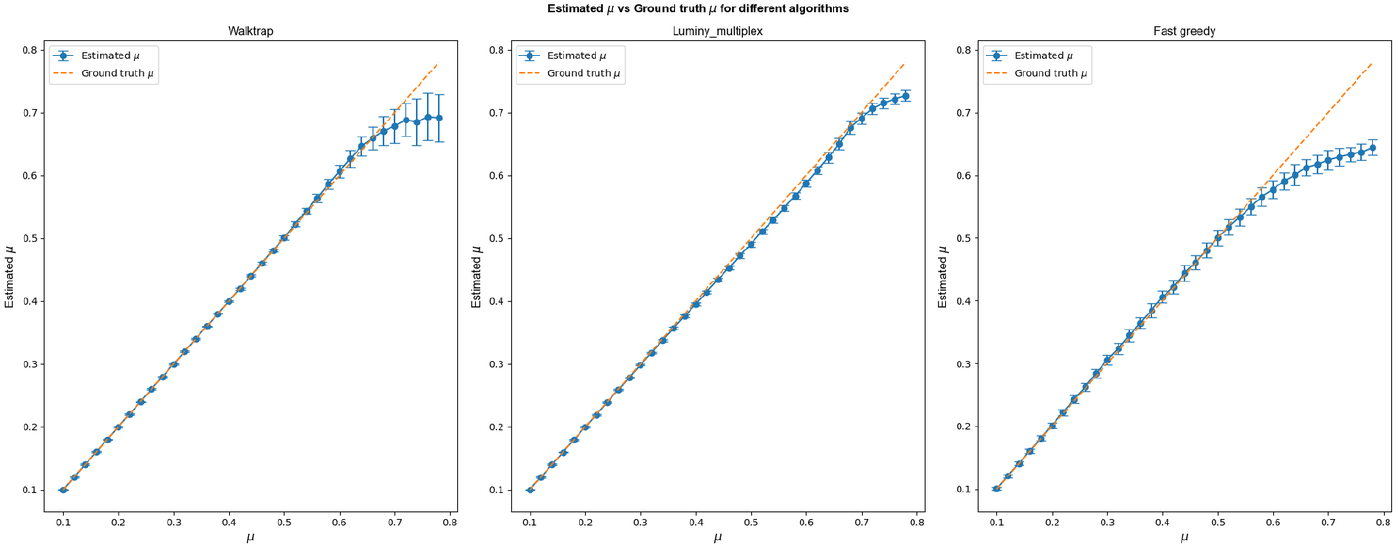
Comparison between the estimated and ground truth mixing parameter (*µ*) The mixing parameter *µ* of the LFR network is varied from 0.1 to 0.8. The plot compares the estimated values of *µ* against the ground truth across different algorithms.

The mixing parameter *µ* of an LFR network varies inversely with modularity of the network, implying that the network is more modular at lower values of *µ* and vice versa, as described by equation 3. In this analysis, we fixed the network size at 1000 nodes throughout. The results for different algorithm categories from the Mixing Parameter-based Analysis are presented in Fig. 2,Fig. 3,Fig. 4,Fig. 5,Fig. 6, arranged into three columns for each result category. The first column shows how Normalized Mutual Information (NMI) changes with increasing mixing parameters for various algorithms. The second column illustrates the cluster ratio, which is defined as the ratio of predicted communities to ground truth. The third column represents the variation in computation time as the mixing parameter increases.

#### 2.1.1 Performance decline follows a sigmoidal pattern

Algorithm performance in different categories, measured by NMI, generally declines at varying rates, typically following a sigmoidal pattern as the mixing parameter *µ* increases from 0.1 to 0.8, showing a sharp decrease at 0.6. This is expected since communities become less defined with higher *µ* values. The best performing algorithms for *µ <* 0.5 and *µ >* 0.5 are summarized in Table 1. In real-world networks, where the mixing parameter *µ* may not be uniform, community detection algorithms can estimate *µ* in unsupervised settings. Fig. 7 compares the estimated values *µ* with the ground truth for Walktrap, Luminy multiplex, and FastGreedy. Luminy multiplex is the most accurate, providing consistently close estimates across all ranges of *µ*, especially in higher mixing scenarios, making it the most reliable algorithm for estimating *µ*.

**Table 1.** Mixing Parameter (*µ*) based Analysis.

| Class | NMI (at lower ($\mu$)) | NMI at higher ($\mu$) | Cluster Ratio | Computation Time | Avg Comp Time (fastest algorithm) |
| --- | --- | --- | --- | --- | --- |
| Stochastic | TripleAHC | MLRMCL | Walktrap | Walktrap | 0.483 sec |
| Modularity Based | Luminy_multiplex | Louvain | Luminy_multiplex | Fastgreedy | 0.039 |
| Hierarchy Based | SpecHier | csbio_iitm-hamming | SpecHier | Dcut | 0.43 |
| Kernel Based | SpecHier | SpecHier | Spectral Clustering | Spectral Clustering | 0.0 |
| Local Search Based | Label Propagation | Shared Neighbor | Label Propagation | Label Propagation | 0.0031 |

Fig. 2,Fig. 3,Fig. 4,Fig. 5,Fig. 6 use error bars to depict the mean result along with the standard deviation (mean ± standard deviation), based on 100 different network realisations at each data point. These error bars provide a visual indication of the variability in the results.

#### 2.1.2 Algorithms with greater granularity maintain stability at high *µ* values

Algorithms that detect communities with finer granularity tend to remain stable even at higher *µ* values. For example, within the category of stochastic methods, TeamCS based on recursive infomap performs well at low *µ* values, but suffers a sharp decline once *µ* exceeds 0.65. In contrast, multi-layer regularized Markov cluster (MLRMCL) shows only moderate performance at low values of *µ*, but maintains stability as *µ* increases. This behavior is further highlighted by the cluster ratio graphs, where MLRMCL in Fig. 2 maintains a cluster ratio close to 4, explaining its moderate performance at lower *µ*. However, its finer community granularity helps sustain its performance at higher *µ* values, a trend also seen in Nextmr in Fig. 5 and shared neighbour in Fig. Fig. 6 clustering methods. Among these, MLRMCL stands out as the best performer in this regard. MLRMCL is particularly suitable when community structures are not well defined and there is no upper limit on the number of predicted communities.

#### 2.1.3 Computation time remains largely unaffected by increasing *µ*

In most cases, the computation time remains consistent as *µ* increases from 0.1 to 0.8. This is evident from the third columns in Fig. 2,Fig. 3,Fig. 4,Fig. 5,Fig. 6,, where most algorithms maintain constant computation times, with few exceptions. A notable outlier is TeamCS in Fig. Fig. 2, which struggles significantly when *µ* exceeds 0.6. This issue arises because TeamCS relies on a recursive sparsification process that breaks down larger clusters into smaller ones until they fall below a certain size threshold, leading to increased computation time as *µ* increases.

### 2.2 Artificial Benchmarks: Network size based Analysis

In this analysis, we have fixed the mixing parameter *µ* at 0.15, while the network size has been varied from 125 to 4000 nodes. The results of different algorithm categories for the analysis based on network size are shown in Fig. 8, Fig. 9,Fig. 10,Fig. 11,Fig. 12 organized into three columns per result category. The first column shows how Normalized Mutual Information (NMI) varies with increasing network size for different algorithms. The second column displays the cluster ratio, which is the ratio of predicted communities to ground truth communities. The third column represents the variation in computation time as the network size increases.

**Fig 8.**
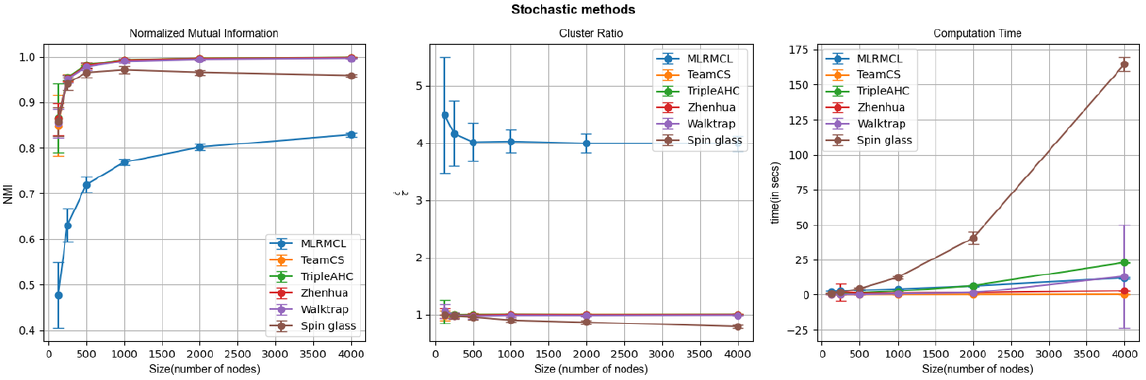
Network size based analysis on Stochastic methods. The size of the LFR network is varied from 125 to 4000 nodes.

**Fig 9.**
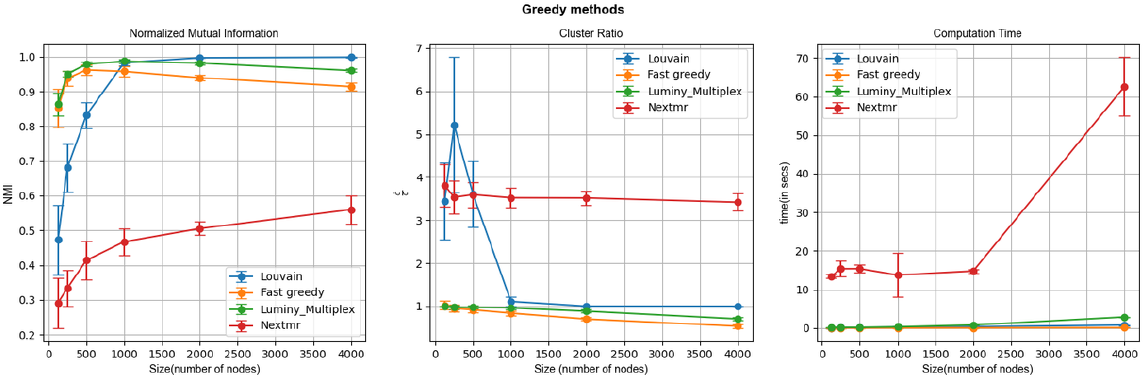
Network size based analysis on Greedy methods. The size of the LFR network is varied from 125 to 4000 nodes.

**Fig 10.**
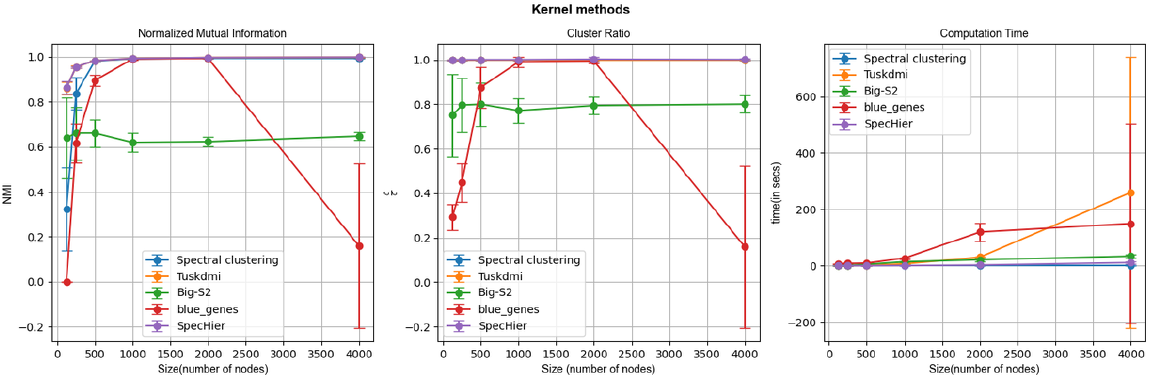
Network size based analysis on Kernel methods. The size of the LFR network is varied from 125 to 4000 nodes.

**Fig 11.**
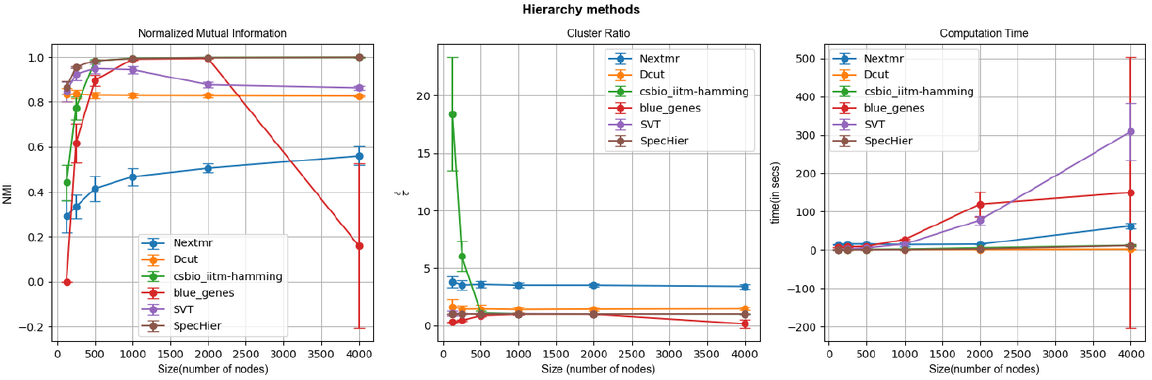
Network size based analysis on Hierarchy methods. The size of the LFR network is varied from 125 to 4000 nodes.

**Fig 12.**
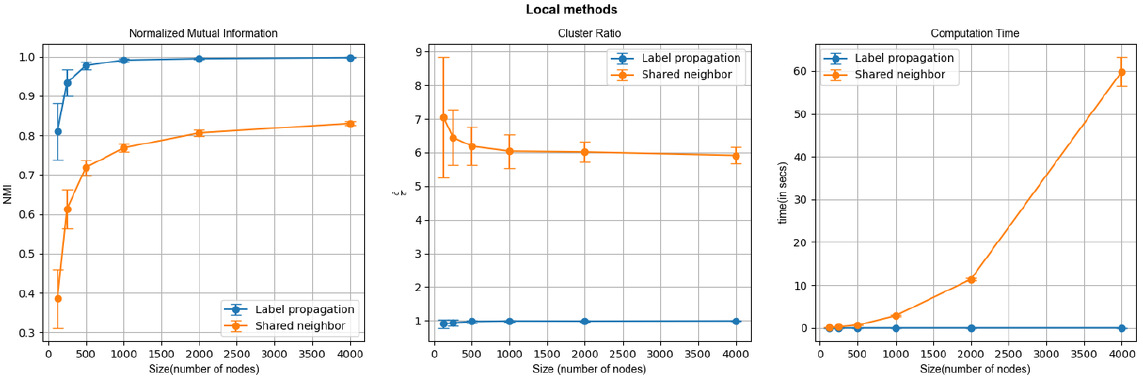
Network size based analysis on Local methods. The size of the LFR network is varied from 125 to 4000 nodes.

Here are the key insights derived from this analysis:

#### 2.2.1 Algorithms underperform with networks smaller than 500 Nodes

Most algorithms across categories experience a sharp increase in NMI as the network size increases from 125 to 500 nodes, after which the performance usually stabilizes. Cluster ratio trends reveal that some algorithms initially deviate at smaller network sizes before stabilizing, while others consistently maintain values above or below 1. This suggests that many algorithms struggle with small networks and only achieve optimal performance once a certain threshold size is reached.

#### 2.2.2 Top-Performing algorithms require time complexity optimization

An interesting observation from the size-based analysis is the behaviour of the computation time as the network size increases from 125 to 4000 nodes. While most algorithms display linear-time complexity with the addition of edges, some demonstrate nonlinear or near-constant complexity. For example, **SpecHier** in Fig. 10 and Fig. 11, a top performer across multiple categories, exhibited non-linear time complexity. In contrast, **Spectral Clustering** in Fig. 10 was the fastest, showing nearly constant time complexity, making it a time-efficient option for larger networks.

The findings of the artificial benchmark analysis are summarized in Table 1 and Table 2.

**Table 2.** Network Size based Analysis.

| Class | NMI (small sized networks) | NMI (large sized networks) | Cluster Ratio | Computation Time | Time Complexity (fastest algorithm) |
| --- | --- | --- | --- | --- | --- |
| Stochastic | TripleAHC,Zhenhua | TripleAHC, TeamCS, Zhenhua | TripleAHC, TeamCS, Zhenhua | TeamCS | $O(n)$ |
| Modularity Based | Multiplex Louvain | Louvain | Multiplex Louvain | Fastgreedy | $O(n)$ |
| Hierarchy Based | SpecHier | SpecHier | SpecHier | Dcut | $O(\sqrt{n})$ |
| Kernel Based | SpecHier | SpecHier | Spectral Clustering | Spectral Clustering | $O(1)$ |
| Local Search Based | Label Propagation | Label Propagation | Label Propagation | Label Propagation | $O(n)$ |

**Table 3.**
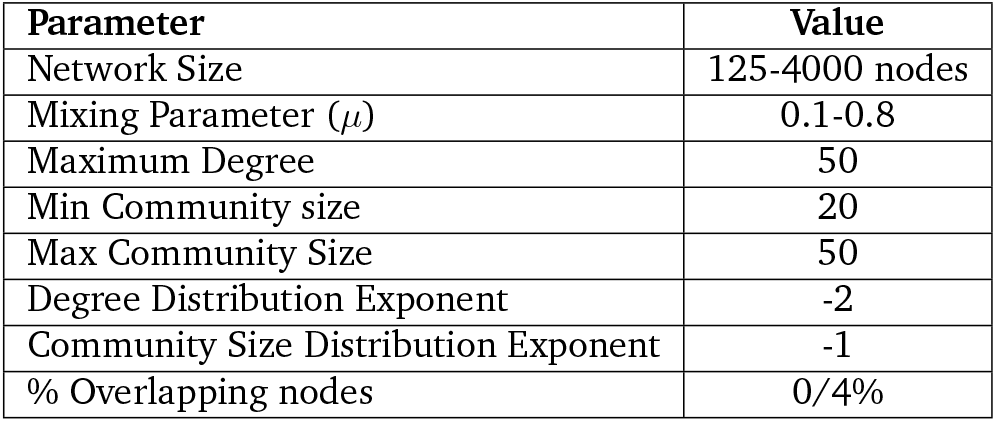
Parameters utilised in the generating LFR Benchmarks. The LFR benchmarks used in the study are undirected weighted networks

| Parameter | Value |
| --- | --- |
| Network Size | 125-4000 nodes |
| Mixing Parameter ($\mu$) | 0.1-0.8 |
| Maximum Degree | 50 |
| Min Community size | 20 |
| Max Community Size | 50 |
| Degree Distribution Exponent | -2 |
| Community Size Distribution Exponent | -1 |
| % Overlapping nodes | 0/4% |

### 2.3 Social Network benchmarking

In the context of social network benchmarking, the manually annotated ground truth did not reveal any consistent trends across metrics such as NMI and modularity.

However, this analysis has been valuable for pinpointing specific bottlenecks where certain algorithms underperform. For instance, the spinglass algorithm, and one of the variants of kernel based algorithms utilising an exp-Laplacian kernel require the network to be a fully connected component for the algorithms to be functional. This insight could not be deduced from the artificial benchmark analysis alone. Visualizations from the social benchmark analysis can be found in the Supplementary Figures S1 Fig, S2 Fig, S3 Fig, S4 Fig and S5 Fig.

### 2.4 Gene Enrichment Analysis

In this subsection, we compare the performance of different algorithms by evaluating the number of statistically significant enrichments (*p*-value ≤ 0.05) identified by these algorithms across communities in the Human and Yeast PPI networks. Additionally, we compare the distribution of community sizes and Fold Enrichment (FE) scores across different algorithms.

#### 2.4.1 Number of statistically significant enrichments per algorithm

The bar charts in Fig. 13 and Fig. 14 show the number of significantly enriched Gene Ontology (GO) terms identified by the algorithms in human and yeast protein-protein interaction (PPI) networks. Here are the key insights:

**Fig 13.**
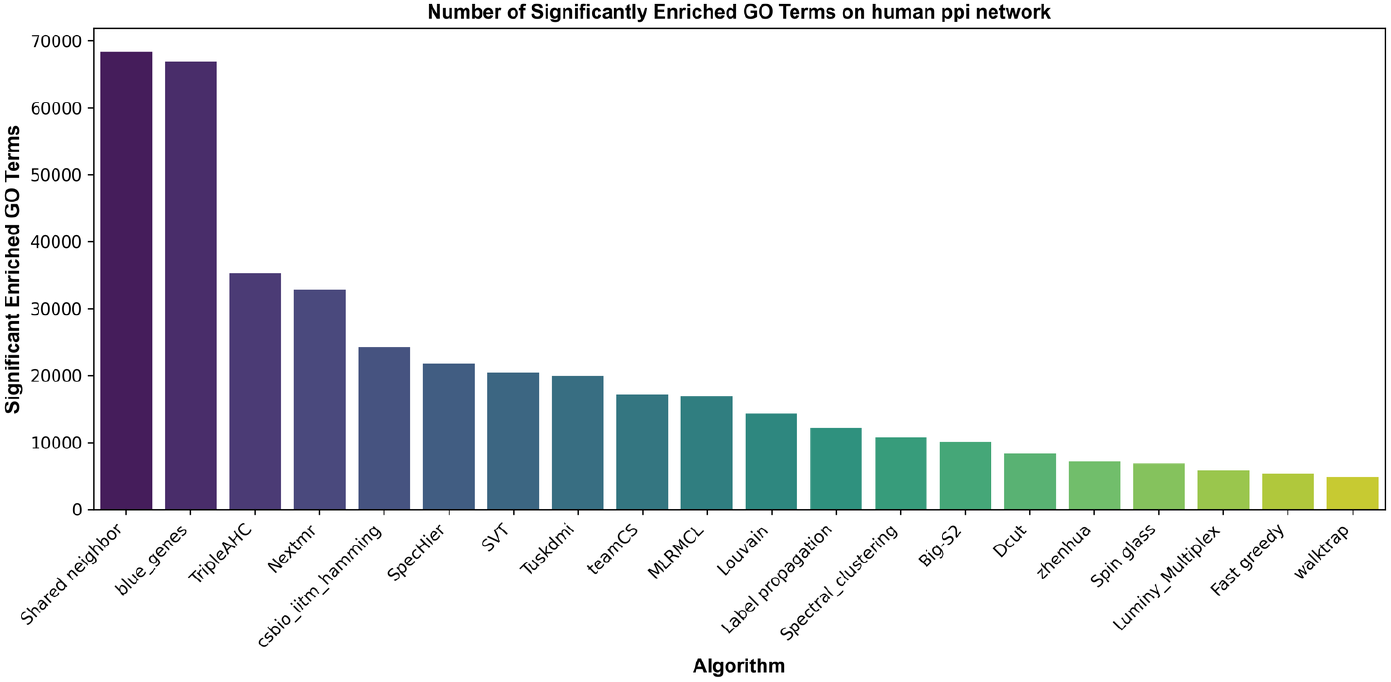
Significant GO Terms on Human PPI. Bar chart depicting the number of statistically significant GO Terms Identified by Algorithms in the Human PPI Network

**Fig 14.**
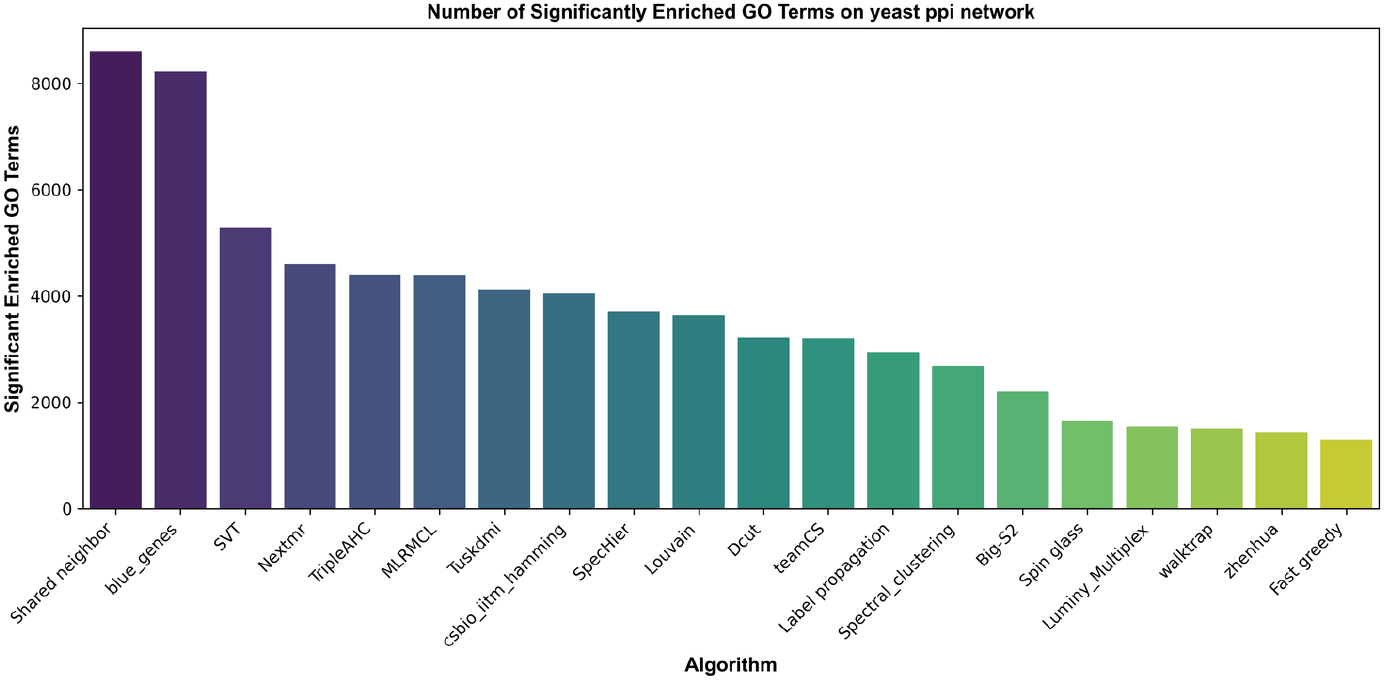
Significant GO Terms on Yeast PPI. Bar chart depicting the number of statistically significant GO Terms Identified by Algorithms in the Yeast PPI Network

##### Human PPI Network

1. **Top Performers**:
  - The **Shared neighbor** and **blue genes** algorithms identify the highest number of significant GO terms ≈ 70, 000, outperforming all others by a significant margin.
  - These algorithms are particularly effective at uncovering enriched functional communities in the human PPI network.
2. **Mid-Performers**: Algorithms like **TripleAHC, NextMR**, and **csbio_iitm-hamming** show moderate performance, identifying between 20,000 to 40,000 significant terms.
3. **Low Performers**: Algorithms such as **walktrap, Fast Greedy**, and **Luminy Multiplex** identify fewer than 10,000 significant terms, indicating limited sensitivity to functional enrichment in human PPI data.

##### Yeast PPI Network

1. **Top Performers**:
  - Similar to the human network, **Shared neighbor and blue_genes** lead in performance, identifying *≈* 8, 000 significant GO terms.
  - The consistent success of these algorithms across both the Human and Yeast PPI networks underscores their robustness and adaptability for functional enrichment analysis.
2. **Mid-Performers**:
  - Algorithms like **SVT, TripleAHC**, and **MLRMCL** demonstrate moderate performance, identifying between 4,000 and 6,000 significant terms.
  - These methods may be suited for smaller or less complex networks.
3. **Low Performers**: Algorithms such as **zhenhua, Fast Greedy**, and **Walktrap** identify the fewest significant GO terms, ranging from 1,000 to 2,000, suggesting limited effectiveness in detecting enriched communities in yeast PPI data.

Based on these results, we recommend using **Shared neighbor** or **blue_genes** for optimal functional enrichment detection. However, both algorithms exhibit slower performance in larger networks. In such cases, **MLRMCL** may serve as a better alternative, offering a reasonable trade-off between speed and enrichment performance.

#### 2.4.2 Comparison of community size distributions and average Fold Enrichment scores

We computed the fold enrichment (FE) scores for Gene Ontology (GO) terms related to Molecular Functions and Biological Processes across the communities detected by each algorithm. The following are the key insights derived from this analysis. The community size distributions are shown in Fig. 15 and Fig. 16, while the average fold enrichment scores are presented in Fig.17 and Fig.18 for the Human and Yeast Protein-Protein Interaction (PPI) networks across different algorithms.

**Fig 15.**
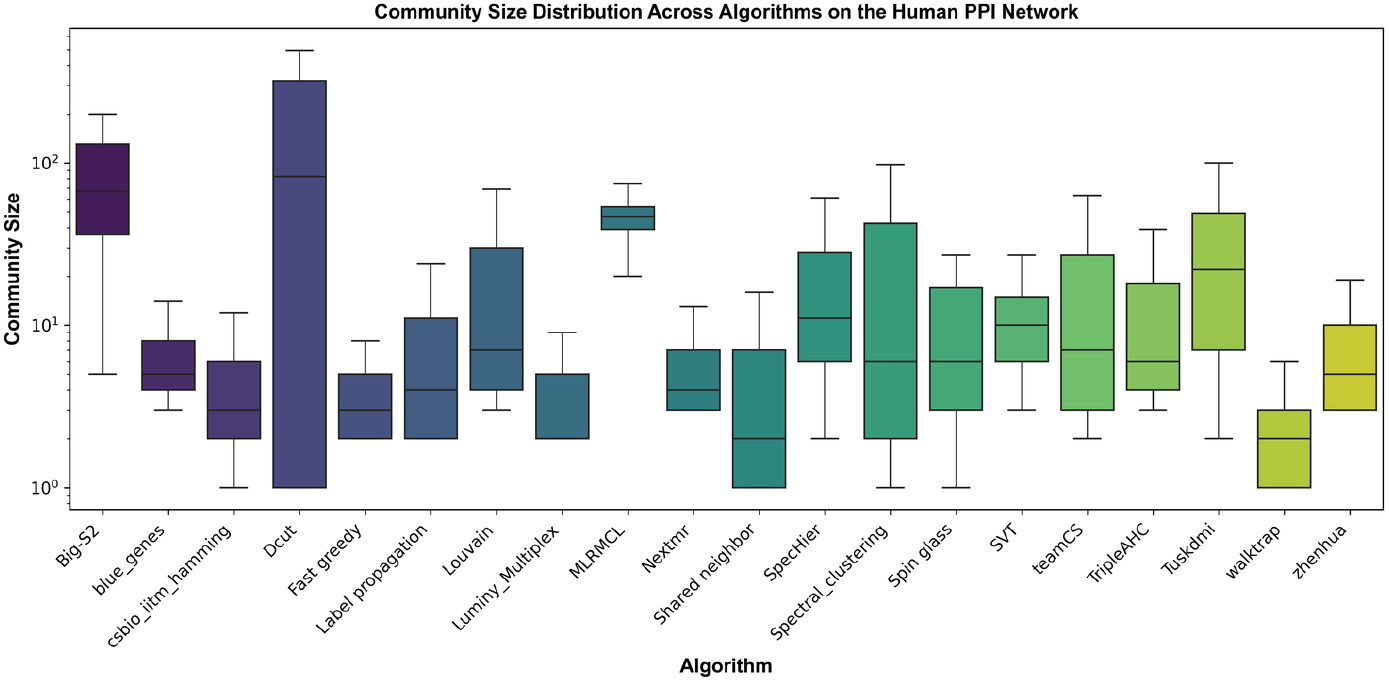
Community Size Distribution in the Human PPI. Box plot depicting the distribution of community sizes in different algorithms in the Human PPI network

**Fig 16.**
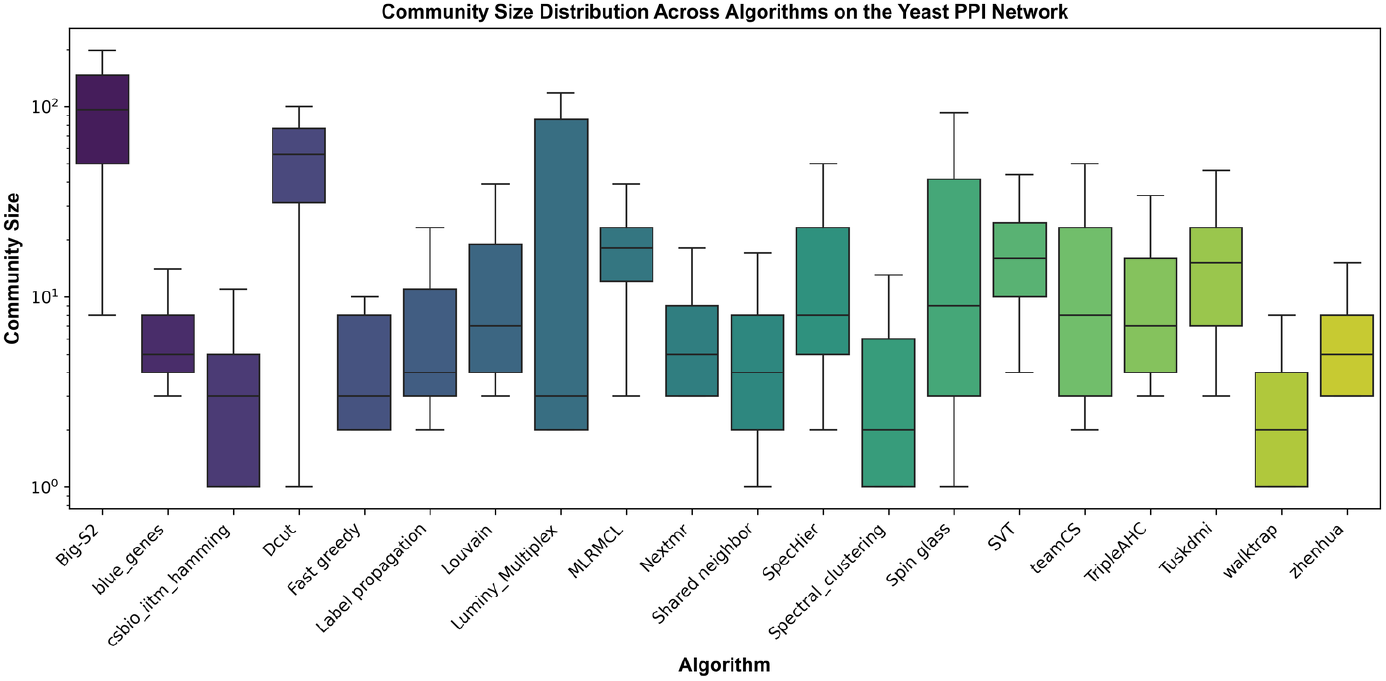
Community Size Distribution in the Yeast PPI. Box plot depicting the distribution of community sizes in different algorithms in the Yeast PPI network

**Fig 17.**
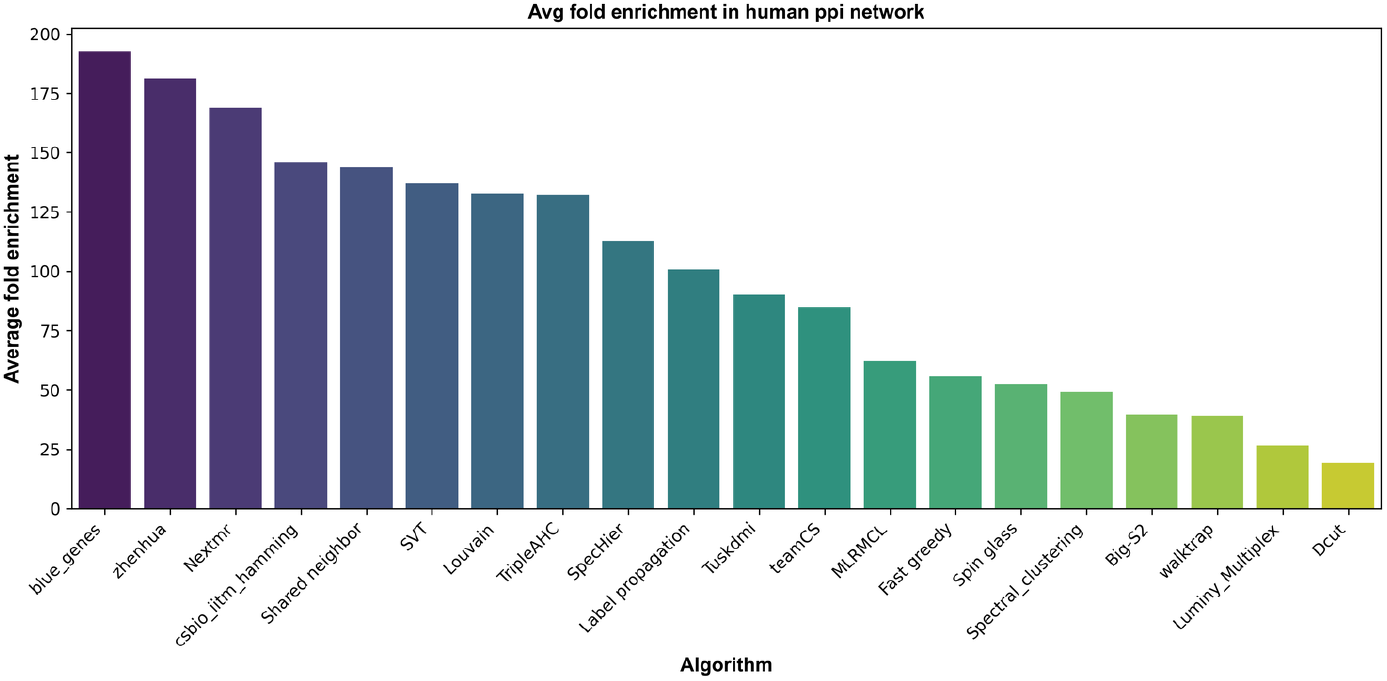
Average Fold Enrichment in the Human PPI. Bar chart depicting the average Fold Enrichment of Go Terms in different algorithms in the Human PPI network

**Fig 18.**
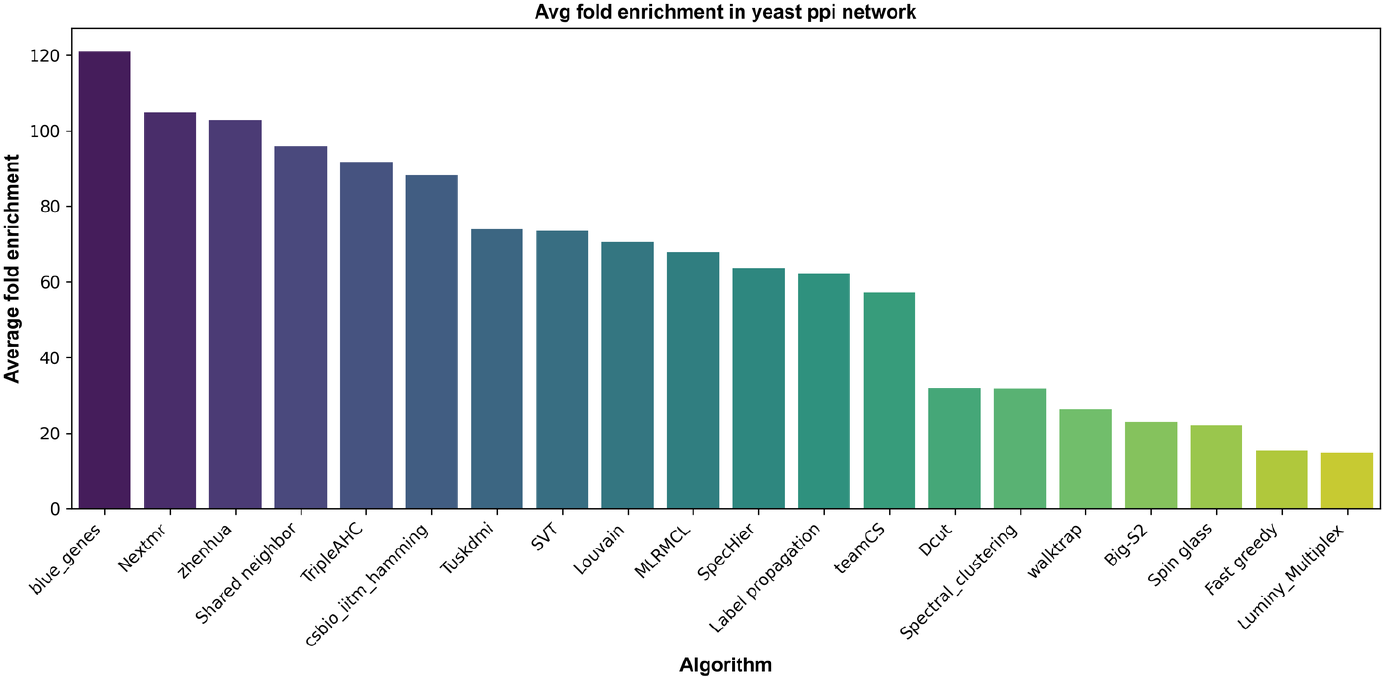
Average Fold Enrichment in the Yeast PPI. Bar chart depicting the average Fold Enrichment of Go Terms in different algorithms in the Yeast PPI network

1. **Top Performing Algorithms** **blue_genes, Zhenhua, Nextmr** stand out as top performers in terms ofaverage fold enrichment across the Human and Yeast PPI networks. These methods typically produce compact community size distributions, suggesting a strong ability to detect smaller, biologically meaningful clusters. Their effectiveness in uncovering functionally relevant communities makes them particularly well-suited for applications like functional annotation and pathway enrichment analysis.
2. **Algorithms with Balanced Performance** **Shared neighbor, Louvain, SVT** exhibit moderate fold enrichment values while maintaining a stable distribution of community sizes. They identify communities of varying sizes without extreme variability or outliers, balancing community size and biological relevance. These algorithms are ideal for general-purpose clustering tasks that require a balance between biological significance and structural diversity. They are applicable in settings where robustness is desired across multiple community scales.
3. **High Variability Algorithms** **Fast greedy, Dcut** are characterized by high variability in community size distribution, often identifying tiny and large clusters. However, their average fold enrichment tends to be lower, suggesting limited biological relevance in functional clustering. These algorithms may be better suited for exploratory analyses or networks with hierarchical structures than biological networks.

##### Recommendations Based on Objectives

Based on the observed performance patterns, we recommend the following algorithms depending on the specific objectives:

a. **For High Biological Relevance**: Algorithms such as *blue_genes, Zhenhua*, or *Nextmr* are recommended due to their high fold enrichment scores and their ability to produce compact biologically significant clusters.
b. **For General-Purpose Clustering**: Balanced methods such as *Louvain, Shared neighbor*, or *SVT* are well-suited for a range of clustering tasks, offering reasonable biological relevance and structural diversity.
c. **For Exploratory Analysis**: High variability algorithms like *Fast greedy* or *Dcut* are recommended for exploratory analyzes, where the goal is to identify diverse clustering patterns rather than optimizing for biological enrichment.

## 3 Methods

### 3.1 Implementation of Community Detection Algorithms

In this work, we implemented a total of twenty-one community detection algorithms, which can be broadly classified into five main types, as elaborated in the rest of this section.

#### 3.1.1 Stochastic Methods

These include methods using key concepts from Infomap, Walktrap, and Spinglass, which rely on probabilistic approaches to detect communities, often using random walks [27, 28] and information theory to identify densely connected clusters within networks.

1. **Infomap:** Infomap is a community detection algorithm that identifies groups of nodes by analysing the flow of random walks through a network. It works by finding a way to describe the paths of these walks as efficiently as possible, revealing communities where the walks tend to stay within certain groups of nodes. The method detects communities by minimizing the overall length of the description, indicating strong internal connections within those groups compared to the rest of the network. The time complexity for this algorithm is *O*(*m*) where *m* denotes the number of edges on the graph.
2. **Walktrap:** The Walktrap algorithm detects communities by simulating random walks within a network, with short walks staying within densely connected nodes. It merges nodes and small clusters based on the frequency of co-visits, progressively building larger communities. The time complexity for this algorithm is *O*(*M* log(*N*)) and for sparse networks is *O*(*N* ^2^ log(*N*)).
3. **Spin glass:** Spin glass is a community detection algorithm inspired by concepts from statistical physics, particularly the behaviour of spin glasses. The network is modeled as a system, where individual nodes represent spins and the edges between them represent interactions. The algorithm aims to minimize an energy function, where communities correspond to low-energy states, implying that nodes within the same community have strong interactions. A unique feature of Spin glass is its simulation of the system’s cooling process (annealing), which helps identify communities as groups of nodes that reach stable, low-energy configurations, indicating strong internal connections. The time complexity of this algorithm is *O*(*N* ^3.2^).

We also implemented some enhanced variants of stochastic Algorithms including:

1. **Multi Layered Markov clustering algorithm:** The Markov Clustering Algorithm (MCL) identifies clusters through iterative expansion and inflation of the adjacency matrix. Multi-Layer Regularized Markov Clustering (MLRMCL) improves MCL by regularizing singleton clusters and using a multilayered approach to reduce computational complexity
2. **TripleAHC:** Walktrap is carried out recursively until the modularity of individual clusters cross a certain threshold
3. **Zhenhua:** Walktrap with further refinement of large clusters using Infomap
4. **TeamCS:** An enhanced version of Infomap with recursive sparsification for larger clusters, utilizing an inverse log-weighted similarity to improve detection accuracy.

#### 3.1.2 Kernel Based Methods

These algorithms mainly utilise concepts of Spectral Clustering and build variations of the same.

##### Spectral clustering

Spectral clustering is a highly effective technique that uses Laplacian’s eigenvalues to reduce the data’s dimensionality before applying a clustering algorithm, such as K-means, to identify clusters. Using the spectral properties of the Laplacian network, spectral clustering can capture complex cluster structures that are not easily identifiable with traditional methods [29]. It works by transforming the data into a lower-dimensional space where clustering can be effectively performed, allowing for improved separation and identification of clusters. The time complexity of this algorithm is *O*(1).

Some of the enhanced variants of kernel-based algorithm we implemented are :

1. **Tuskdmi:** Applying spectral clustering to a custom Diffusion State Distance (DSD) matrix
2. **SpecHier:** Using spectral clustering to determine initial clusters, followed by denoising and applying agglomerative clustering on larger subgraphs
3. **Big-S2:** Performing spectral clustering followed by iterative refinement to split larger communities into sub-communities
4. **blue genes:** Using spectral clustering on Laplacian exponential diffusion kernels to capture complex community structures

#### 3.1.3 Modularity based Methods

Modularity-based algorithms are designed to identify community structures in networks by optimizing a quality measure known as modularity [30, 31]. Modularity quantifies the strength of the division of a network into communities, with higher values indicating a more pronounced community structure. These algorithms work by partitioning the network to maximize modularity, which measures the density of edges within communities relative to edges between communities.

Common approaches include Louvain and Fast Greedy algorithms, which iteratively refine community assignments to improve modularity scores.

1. **Fast greedy:** Fast Greedy performs hierarchical agglomerative clustering by merging communities to maximize the gain in modularity at each step.
2. **Louvain:** Louvain employs a multilevel approach where nodes are iteratively moved between communities to identify shifts that lead to the most significant increase in modularity. After forming communities, nodes within each community are aggregated into supernodes, and the process is repeated recursively.
3. **Luminy_multiplex:** We implemented a variation of the Louvain method, which introduces randomization and the ability to detect communities in different layers of the network.

#### 3.1.4 Hierarchy based Algorithms

The Girvan-Newman algorithm is a classic community detection method based on the concepts of divisive hierarchical clustering. It iteratively removes edges with the highest betweenness centrality, splitting the network into smaller communities. Unlike other methods that group similar nodes, Girvan-Newman reveals communities by breaking key connections between them. Although effective for small-medium sized networks, it can be computationally intensive for larger networks with a time complexity of O(E^2^N). However, due to its primary design for unweighted networks, it has not been included in Figs. Fig. 2,Fig. 3,Fig. 4,Fig. 5,Fig. 6 for the weighted LFR benchmark analysis.

Most hierarchy-based clustering algorithms mainly use the concepts of agglomerative clustering [32]. These algorithms start with individual nodes as separate clusters and iteratively merge the closest clusters, updating the distance matrix at each step to build a hierarchical structure, often visualized as a dendrogram.

We implemented some variations of hierarchical clustering including:

1. **csbio_iitm-hamming:** In this method, an ensemble matrix is constructed from partitions at different resolution levels and hierarchical clustering is performed based on the frequency with which nodes appear together in the same community across resolutions [33].
2. **Nextmr:** Here, A Topology Overlap Matrix (TOM) is constructed from the graph’s adjacency matrix and hierarchical clustering is applied followed by modularity and entropy-based refinement.
3. **Dcut:** In this method, hierarchical clustering is performed on the similarity matrix of the graph (e.g., Jaccard similarity) to build a tree, and final clusters are obtained through optimal recursive splitting of the tree [34].
4. **Singular Value Thresholding (SVT):** This method utilises a latent feature space representation of the network using Singular Value Thresholding, where agglomerative and recursive clustering is performed.

#### 3.1.5 Local Search based Algorithms

Local search community detection algorithms cluster nodes by focusing on local patterns within the network instead of optimising a global metric like modularity. This approach is particularly efficient for large networks, where local interactions are the key to uncovering meaningful communities.

##### Label propagation

This method iteratively updates node labels by assigning each node the most frequent label found in its neighborhood, continuing the process until the community structure stabilizes. The time complexity of this algorithm is Õ(m)

##### Shared neighbor Clustering

This approach iteratively merges nodes by focusing on their degree and connectivity, particularly emphasizing integrating clusters with fewer nodes to enhance community cohesion.

### 3.2 Benchmarking Strategies

We benchmarked the community detection algorithms described above using both social networks and artificial benchmarks with known ground-truth communities. For social network analysis, we used eight different real-world networks with known communities. However, in social networks, the concept of a ground-truth community is often ambiguous and open to interpretation, which limits the reliability of such studies.

Consequently, the number of studies that employ social networks to benchmark community detection algorithms remains limited.

For artificial benchmark analysis, we used weighted and undirected LFR networks. One of the first synthetic benchmarks, still widely used today, is the Girvan–Newman benchmark [35]. However, the Girvan-Newman benchmark presents a very simplistic network structure, where all nodes have the same degree, and all communities are of standard size. To address these limitations, the LFR benchmarks introduced by

Lancichinetti *et al*. [36], offering a more realistic approach by generating artificial networks that account for node degree and community size heterogeneity, both of which follow a power-law distribution.

Furthermore, the generalized LFR benchmarks [37] represent another advance, incorporating features such as orphan nodes and allowing different mixing parameters within the same network, making them even more suitable for complex network analysis.

We implemented our community detection algorithms in a semi-supervised setting, where the necessary parameters for each algorithm, such as community length and size limits, were tuned based on the available ground truth data. The performance of the algorithms was evaluated primarily using the normalized mutual information metric (NMI) to measure the similarity between the output clusters and the ground truth.

In addition to NMI, we further analyzed the granularity of the clusters by examining the cluster ratio, which provides insight into the distribution of cluster sizes. Moreover, we conducted an analysis of the computational efficiency of different algorithms by evaluating their time complexity [19].

#### 3.2.1 Metrics Used

- **Normalized Mutual Information:** The Normalized Mutual Information (NMI) is a standard metric borrowed from the concepts of Information Theory [38]. NMI comparing two community partitions is given by the following formula:

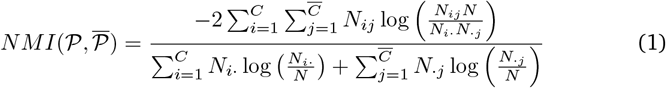

For the calculation of NMI, a confusion matrix **N** is derived first where an element in the matrix *N*_*i,j*_ is obtained as the number of nodes in *j*^th^ predicted communities found in *i*^th^ real community. *N* refers to the number of nodes in the network. *N*_*i*._ refers to the number of nodes in *i*^th^ real community and *N*_.*j*_ refers to the number of nodes in *j*^th^ predicted community. NMI between the predicted and real community partitions scales between 0 and 1, indicating no overlap and perfect partition, respectively.
- **Cluster Ratio:** Cluster Ratio is simply the ratio of the number of predicted communities to the number of communities in the ground truth partitions. Ideally expected to be 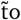 1, the cluster ratio indicates the granularity with which the algorithms detect communities.
- **Computation time:** Computation time refers to the runtime duration of different algorithms for a network of a given size.
- **Modularity:** Modularity quantifies the strength of the division of a network into communities, with higher values indicating a more pronounced community structure [31]. The modularity *Q* for a given partition is given by

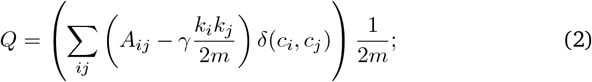

where A is the adjacency matrix of the network, *k*_i_ and *k*_j_ are the degrees of nodes *i* and *j*, and *m* is the total number of edges in the network. *γ* is a resolution parameter. *δ*(*c*_i,_, *c*_j_) is the Kronecker delta function, which takes the value 1 if *i* and *j* are in the same community, and 0, otherwise. This metric can be used to score community partitions by a given algorithm when the ground truth partition is not available.

In summary, we used the Normalized Mutual Information score as the primary metric to score algorithms for their performance in different network structures. In addition, the cluster ratio and computation time serve as supporting metrics that provide additional insights. We used modularity as a metric primarily in social network benchmarking, where the annotation of ground-truth communities is not very clear.

### 3.3 Artificial Benchmarks

In case of artificial benchmarks, we performed two types of analyses using the LFR class of networks, by varying Mixing parameter(*µ*), and also by varying the size of the network, as described in the following sections:

#### 3.3.1 Varying Mixing Parameter (*µ*)

The mixing parameter *µ* for a given network is calculated using the following equation:

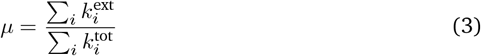

where 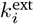 represents the external degree of node i, i.e. the number of connections that node i has outside its community, and 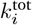 represents the total degree of node i, which is the total number of connections the node has. A lower value of the mixing parameter *µ* indicates that the network is more modular, meaning that the nodes tend to have more connections within their community than outside it.

In this analysis, we varied the mixing parameter of the system from 0.1 to 0.8 generating 100 different realisations of the weighted, undirected network of 1000 nodes at each value of *µ* to derive statistically significant results.

#### 3.3.2 Varying Network size

In this analysis, we fixed the mixing parameter (*µ*) of the network at 0.15 with 4% of overlapping nodes, while the network size varies from 125 to 4000 nodes. For each network size, we generated 100 different realizations of undirected weighted networks. This approach is particularly useful for evaluating the time complexity of various algorithms.

#### 3.3.3 Social Networks

Numerous social networks with annotated ground truth data are available in various online network repositories. Of the eight networks that we chose for this analysis, five are undirected and three are directed, all of which are unweighted. Table 4 and Table 4 provide a summary of the networks used in the study.

**Table 4.** Social networks. Eight popular social networks have been employed in the study

| Network | Nature | Clusters | Nodes | Edges |
| --- | --- | --- | --- | --- |
| Adjnoun | Undirected | 2 | 112 | 425 |
| Dolphins | Undirected | 4 | 62 | 159 |
| Football | Undirected | 12 | 114 | 613 |
| Polbooks | Undirected | 3 | 105 | 441 |
| Polblogs | Directed | 2 | 1,490 | 19,090 |
| Karate | Undirected | 2 | 34 | 64 |
| Email | Directed | 40 | 1,000 | 25,571 |
| Cora | Directed | 7 | 2,708 | 5,429 |

**Table 5.** Biological Networks. PPI network configurations

| Organism | Threshold | Nodes | Edges |
| --- | --- | --- | --- |
| Human | 999 | 12,323 | 28,284 |
| Human | 900 | 4,939 | 201,712 |
| Human | 700 | 16,201 | 473,860 |
| Yeast | 700 | 5,791 | 208,376 |

### 3.4 Gene Enrichment Analysis

In addition to the benchmark analysis conducted on artificial and social networks, we performed biological benchmarking using human and yeast protein-protein interaction (PPI) networks obtained from the STRING database. We applied a threshold for filtering edges for most algorithms based on a combined score of ≥ 700, resulting in 16,201 nodes and 473,860 edges in the Human PPI network and 5,791 nodes with 208,376 edges in the Yeast PPI network. Both networks are undirected and weighted, respectively.

However, for the blue genes and SVT algorithms applied to the Human PPI network, we set higher thresholds of 900 and 999, respectively, to accommodate memory and time complexity constraints. These adjustments resulted in a Human PPI network consisting of 12,323 nodes and 201,712 edges for the blue genes algorithm and 4,939 nodes with 28,284 edges for the SVT algorithm.

Since ground truth communities were unavailable for these networks, we conducted a gene enrichment analysis using Gene Ontology (GO) term annotations, focusing specifically on GO terms associated with Molecular Function and Biological Process.

To prepare the data, we first mapped the STRING accession IDs to UniProt IDs based on their annotations in the GO database. Following this preprocessing step, we computed each algorithm’s fold enrichment (FE) score across different communities and GO terms. The FE score is defined as:

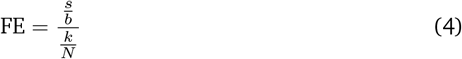

where *s* represents the number of proteins common to both the community and the proteins associated with the GO term, *b* denotes the total number of proteins related to the GO term, *k* is the number of proteins within the community and *N* is the total number of proteins in the selected PPI network.

### 3.5 Implementation

The algorithms implemented in this work were originally developed in Python, R, Matlab, and Java. We have rewritten the BigS2 algorithm, initially written in Matlab, in Python3. We have integrated all algorithms into a single Python script that serves as the main interface. We carried out the benchmark analysis in Python3. We created Fig. 1 using Canva, while generating the remaining figures were using Python3. We conducted all benchmark experiments on a system with an Intel64 Family 6 Model 94 processor (2.71 GHz), 16 GB of RAM, and a 512 GB HDD. The system runs Microsoft Windows 10 Pro, version 10.0.19045. The system was equipped with WSL2 (Windows Subsystem for Linux 2), which allows the use of Linux-based tools and scripts alongside Windows applications.

## 5 Discussion

In this study, we present a comprehensive implementation and benchmark analysis of both classical and hybrid classical algorithms for community detection. Our evaluation scheme spans three categories of networks: artificial, social, and biological, where we offer unique insights into algorithm performance.

Since the ground truth community structures are known in artificial networks, this allowed for direct quantitative evaluation using the Normalized Mutual Information (NMI) metric. The results revealed that the algorithm performance often follows a sigmoidal or linear degradation pattern as the mixing parameter (*µ*) increases from 0 to 1, reflecting the increasing difficulty in distinguishing community boundaries.

The artificial benchmarks described above may not fully capture the mesoscale and microscale properties of real-life networks. The work of Xiao *et al*. [39], which constructs benchmarks by rewiring real-world networks while preserving their key properties, could address the limitations of current real and artificial benchmarks. Although the notion of ground-truth communities in social networks is not very clear, the benchmark analysis highlighted practical limitations and bottlenecks in certain algorithms. For example, the Spinglass algorithm requires a fully connected network, which is often unrealistic for large-scale sparse social graphs. This analysis informed us about algorithmic constraints in real-world applications.

Biological networks enabled the assessment of algorithms based on their ability to uncover biologically enriched communities, modules that are functionally relevant in molecular systems. Among the evaluated methods, the Blue Genes algorithm, which leverages an exponential Laplacian kernel, and Zhenhua’s method, which combines Walktrap clustering with further refinement using Infomap, emerged as top performers in this domain.

Algorithms like SpecHier and TripleAHC, which excelled in artificial benchmark analysis, performed only moderately in biological networks. Conversely, algorithms such as NextMR and Shared neighbor, which performed less impressively in artificial benchmark analysis, achieved high fold enrichment scores in biological benchmarking. These observations yet again highlight the fundamental difference between power-law networks and biological networks.

A critical dimension which would require optimization in future work is the management of computational resources. Both the Shared neighbor and blue genes algorithms exhibit non-linear increases in computation time as network size grows, creating significant challenges during inference. Similarly, SVT’s reliance on Singular

Value Decomposition through scipy imposes memory constraints due to the intensive matrix operations involved. While the classical algorithms we implemented offer limited scope for GPU optimizations, future research could explore deep learning techniques such as graph neural networks, graph GANs, and graph autoencoders to achieve better computational efficiency.

The field is increasingly headed toward overlapping community detection algorithms. Benchmarking these methods could significantly advance ongoing research efforts.

Fortunato and Lancichinetti have developed hierarchical LFR benchmarks for weighted directed networks with overlapping communities, providing valuable tools for evaluation. Extending our analysis to evolving networks, particularly those with overlapping communities, would offer deeper insights into dynamic network structures. Additionally, Fortunato *et al*. have designed stochastic block model benchmarks specifically for dynamically evolving networks [40].

In summary, we present a comprehensive benchmarking of community detection algorithms, emphasizing the importance of context-aware evaluation. By integrating diverse network types, reproducible tools, and biological validation, we reveal key algorithmic trade-offs and domain-specific behaviors. Our findings underscore that performance on synthetic benchmarks does not necessarily translate to real-world utility, particularly in biological networks, guiding more informed and context-sensitive algorithm selection.

## Supporting information

**S1 Fig. Social Network Benchmarking on Stochastic Methods:** Based on NMI, Cluster ratio, Computation time, Modularity

**S2 Fig. Social Network Benchmarking on Kernel based Methods:** Based on NMI, Cluster ratio, Computation time, Modularity

**S3 Fig. Social Network Benchmarking on Modularity based Methods** Based on NMI, Cluster ratio, Computation time, Modularity

**S4 Fig. Social Network Benchmarking on Hierarchy based Methods** Based on NMI, Cluster ratio, Computation time, Modularity

**S5 Fig. Social Network Benchmarking on Local Search based Methods** Based on NMI, Cluster ratio, Computation time, Modularity

**S6 Fig. Significant Terms Vs Average Fold Enrichment in the Human PPI network** Scatter plot depicting significant Terms Vs Average Fold Enrichment for the 20 algorithms in the human PPI Network

**S7 Fig. Significant Terms Vs Average Fold Enrichment in the Yeast PPI network** Scatter plot depicting significant Terms Vs Average Fold Enrichment for the 20 algorithms in the yeast PPI Network

**S8 Appendix. Gene Enrichment Analysis Summary for the Human PPI network** CSV File Summarizing the Gene Enrichment Analysis results from different algorithms on the Human PPI Network

**S9 Appendix. Gene Enrichment Analysis Summary for the Yeast PPI network** CSV File Summarizing the Gene Enrichment Analysis results from different algorithms on the Yeast PPI Network

## Acknowledgments

The authors acknowledge the support of the PG Senapathy Center for Computing Resources for providing a High Performance Computing Environment.

## Data availability

The social networks used for the benchmarking in this analysis were sourced from **Network Repository**, [41]. Furthermore, artificial LFR benchmarks were generated by compiling the C++ codes for weighted and undirected networks available on Santo Fortunato’s official website [42] link.

## Code availability

All algorithms used are interfaced with one Python main script, and a complete Python package has been created. The Python package can be cloned and installed locally in the system from this repository. In addition, the module can be imported into an external script or can be accessed through the command-line interface. The Python module takes in an edgelist file or a networkx graph as input and returns a list of lists containing the communities and optionally writes the output into a.txt file when the path for the same is provided. An example of usage utilising the Python command-line interface is highlighted below.

~~~
Python -m community_detection_1.main network.dat
MLRMCL --output_file output.txt
--algorithm_args largest=50
~~~

Here, community_detection_1 is the name of the Python package. The input file is network.dat and the algorithm chosen for this above example is MLRMCL. In addition, the output path is specified as output.txt and largest=50 is one of the algorithm parameters.

