## Supplementary figures and images for "Extensive Benchmarking of Community Detection Algorithms"

### S1_Fig

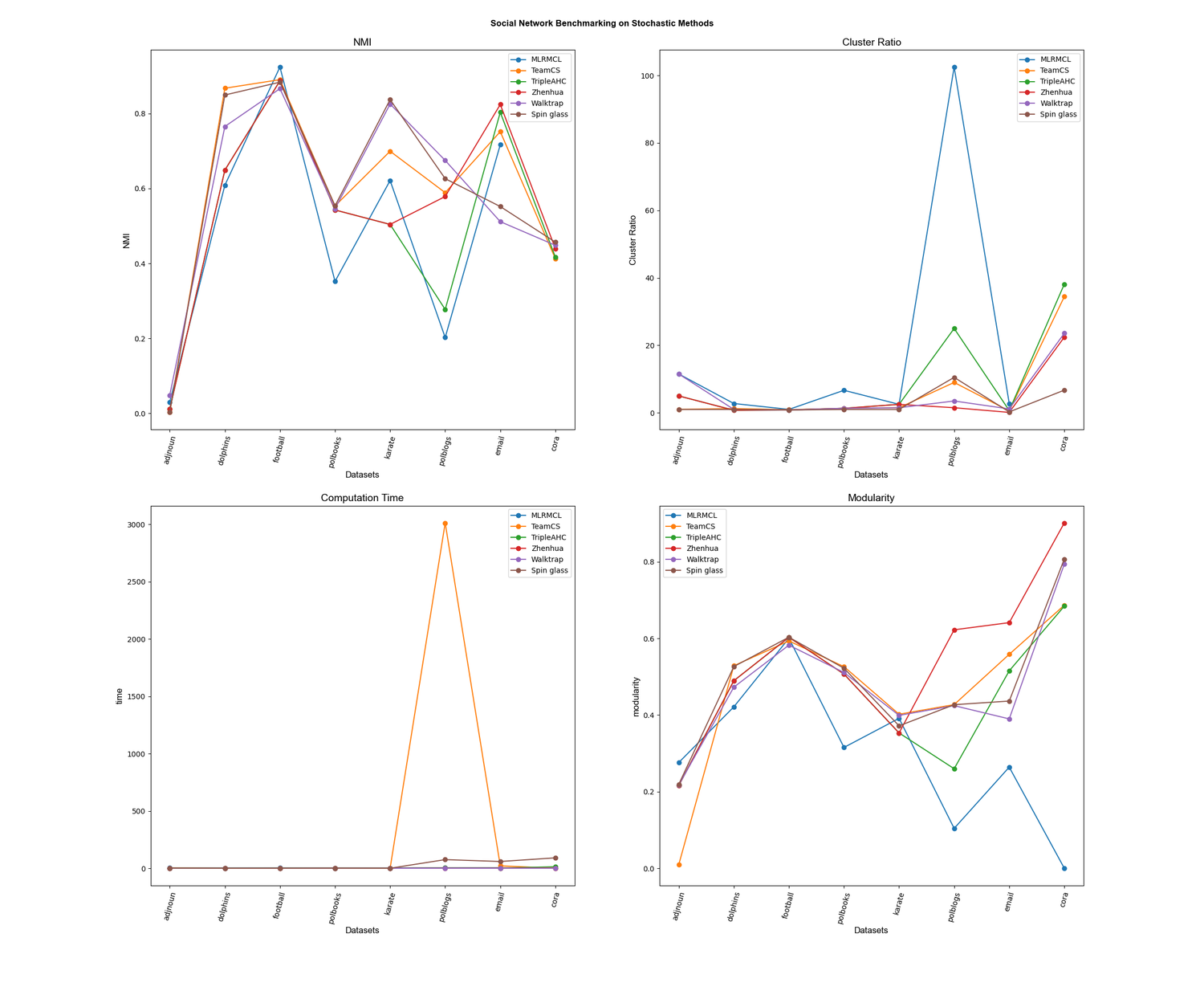

### S2_Fig

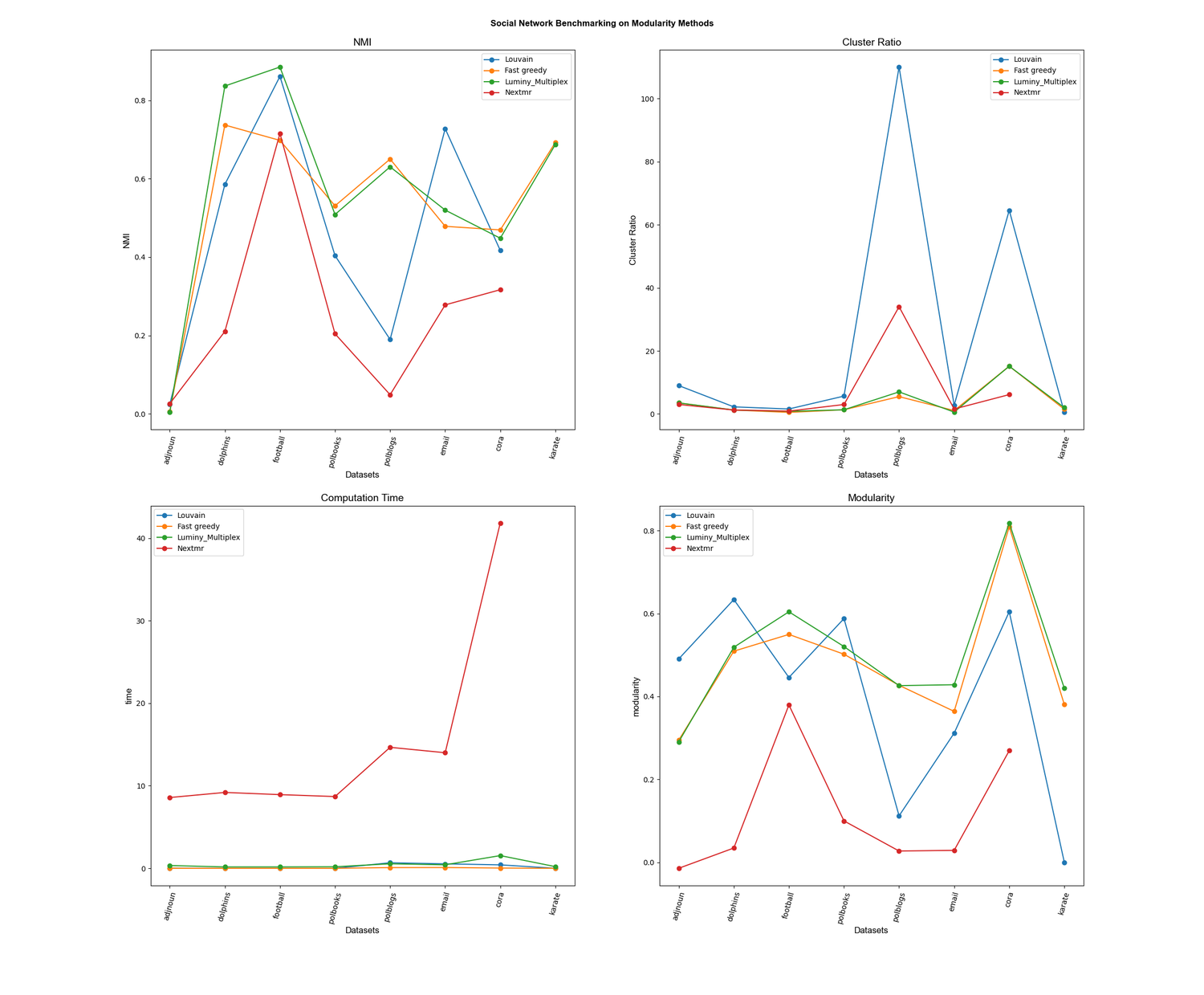

### S3_Fig

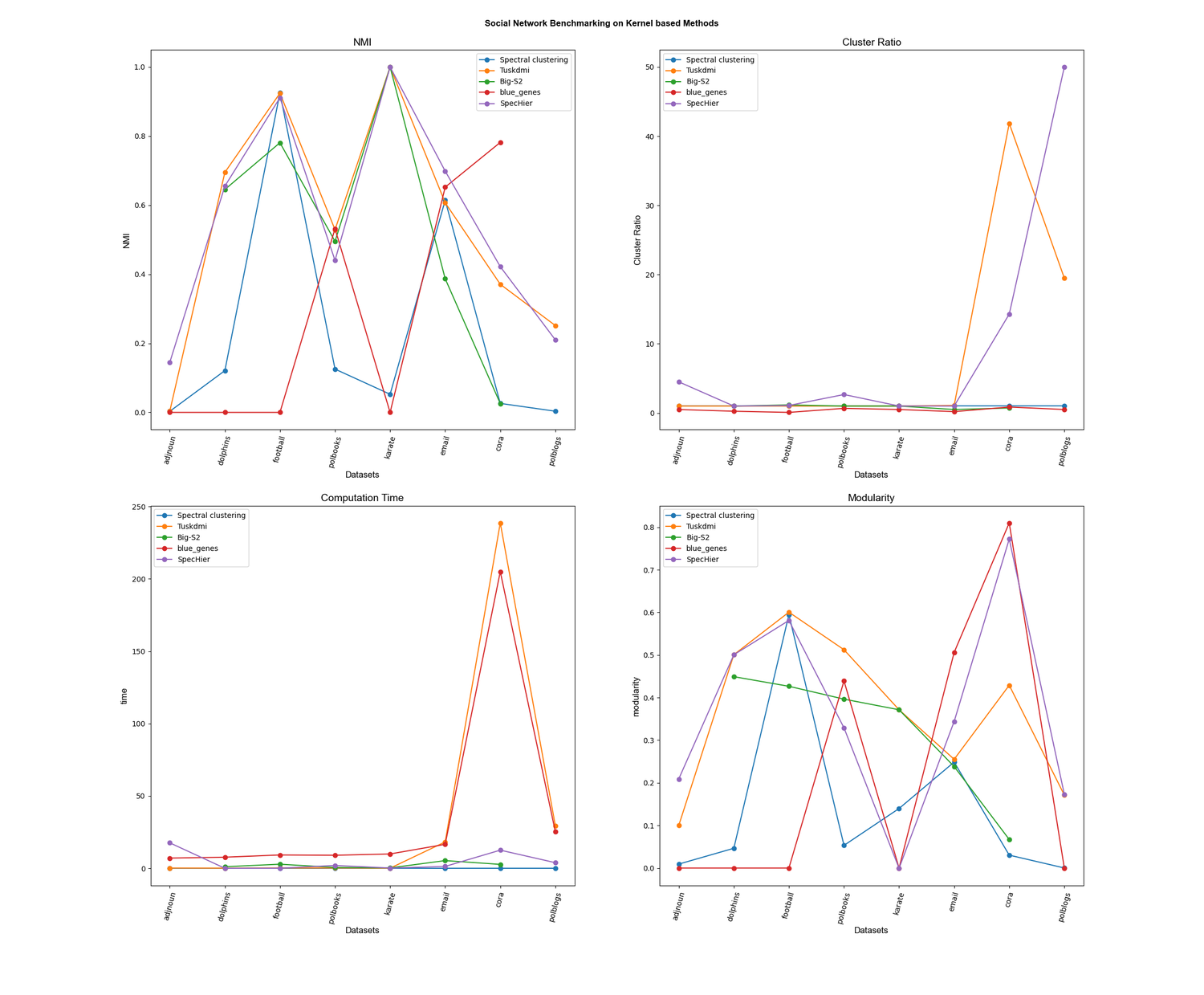

### S4_Fig

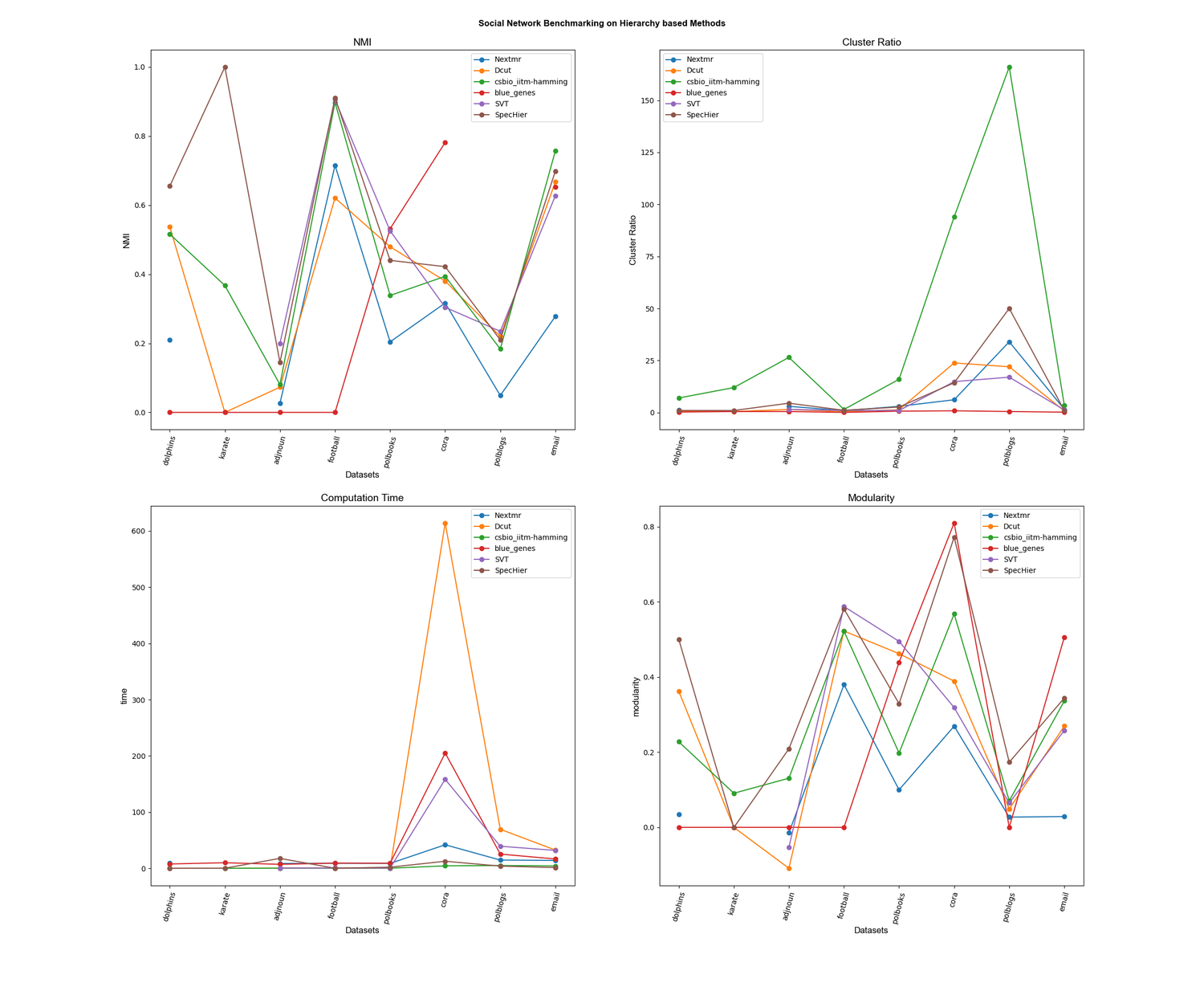

### S5_Fig

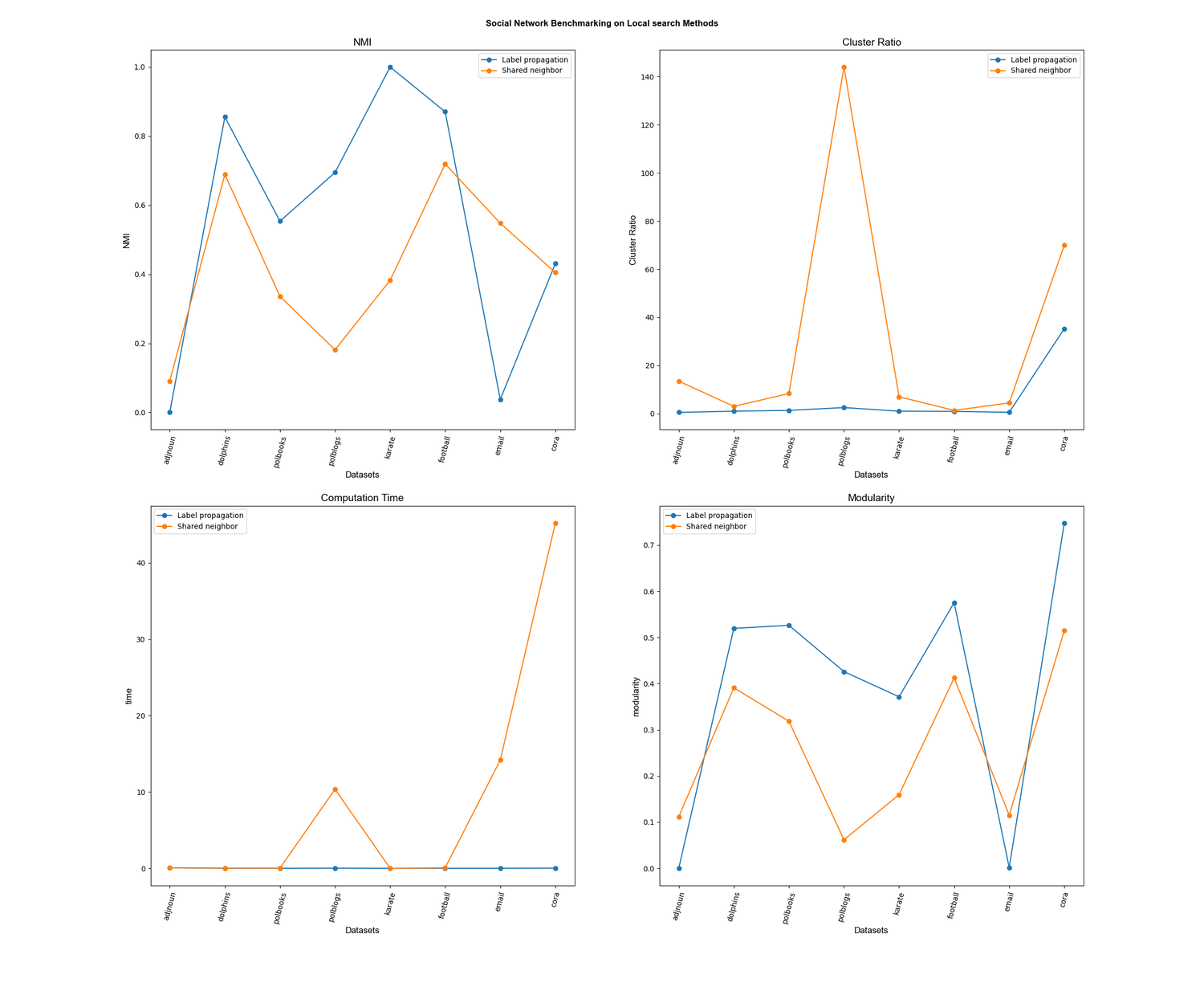

### S6_Fig

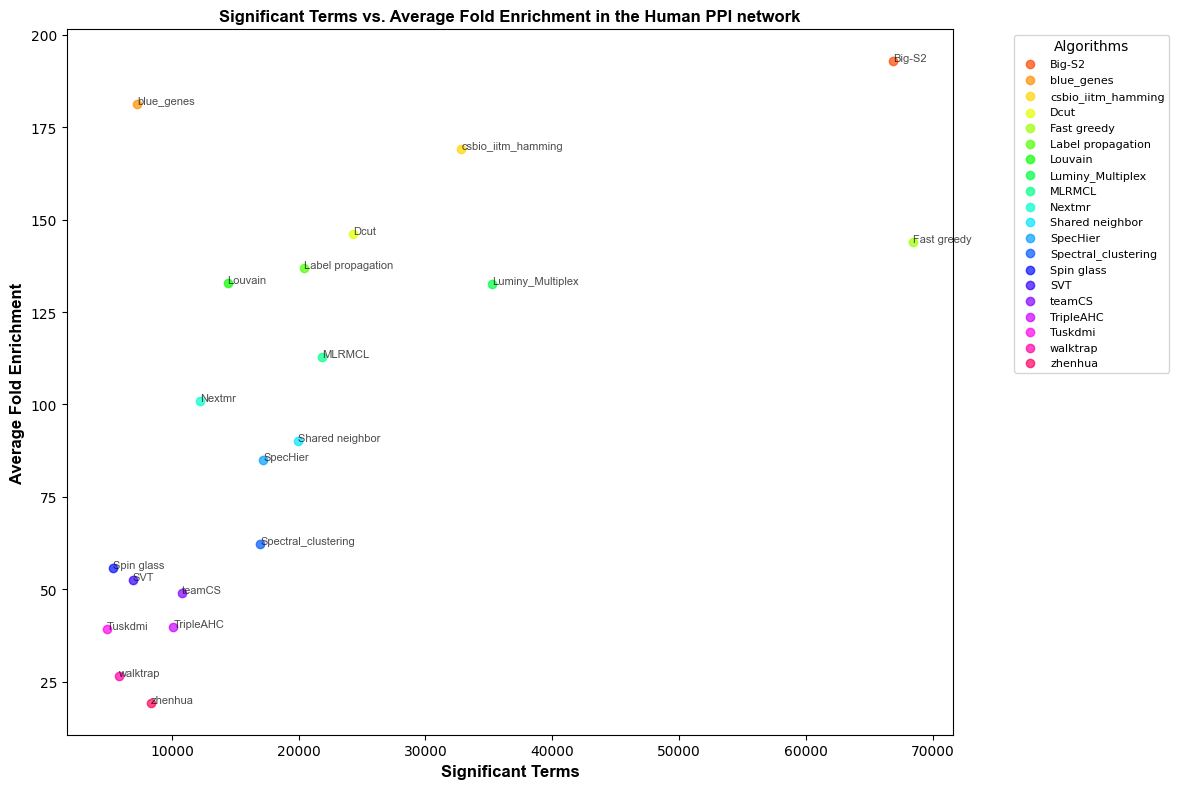

### S7_Fig

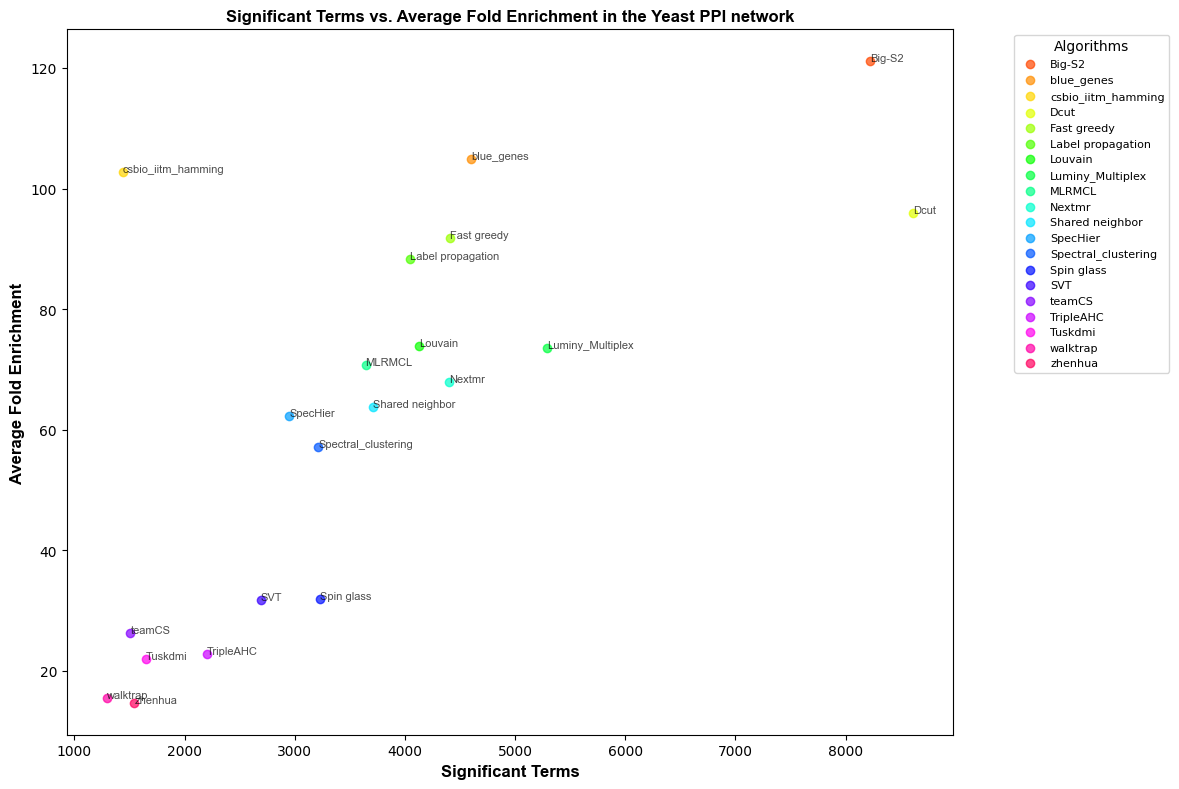
